# An integrated bioinformatics approach to identify candidate biomarkers and the evaluation of drugs for pheochromocytoma

**DOI:** 10.1101/2023.02.17.529021

**Authors:** S Prashanth Javali, J Jeyakanthan, Chun-Jung Chen, Mohanapriya Arumugam

**Affiliations:** Department of Biomedical Genetics, School of Biosciences and Technology, Vellore Institute of Technology, Vellore, Tamilnadu, India; Department of Bioinformatics, Alagappa University, Karaikudi, Tamil Nadu, India; Laboratory of Protein Crystallography and Structural Biology, National Synchrotron Radiation Research Center (NSRRC) No. 101, Hsin-Ann Road, Hsinchu, TAIWAN; Department of Biotechnology, Vellore Institute of Technology, Vellore, Tamilnadu, India

## Abstract

**Objectives:** Identification of the most potential druggable targets and drugs for treating pheochromocytoma by understanding the molecular mechanisms in the progression of pheochromocytoma and discovering additional biomarkers for early diagnosis.

**Methods:** The differential gene expression in microarray data was analyzed using CRAN packages such as Geoquery, Affy, Limma, Tidyverse, GOchord, etc., and functional and pathway enrichment was performed through the David database, Further PPI network is constructed by retrieving the Protein partners were retrieved from the STRING database. The top 3 ranked gene products from cytohubba were docked with the ligand molecules, which are all in phase III trials. The complex with the highest binding affinity was simulated using molecular dynamics simulations for 100ns, PCA and FEL analysis were performed to determine the stability of the complex at each time interval additionally miRNA and TRFs were predicted for the prominent genes.

**Results:** As a result of functional enrichment, most of the 97 differentially expressed genes (DEGs) are involved in cancer-causing pathways. Pheochromocytomas were associated with CDC42, VEGFA, PIK3R1, ITGAV, and LAMB-1, among prominent genes among the ten hub genes. Simulated results verified the effectiveness of targeting VEGFA with Lenvatinib.

**Conclusions:** We found that differentially expressed genes in the interactome and hub genes with high correlations could serve as biomarkers, along with their miRNAs and TRFs. By targeting VEGFA as a drug target, Lenvatinib displays a high affinity for VEGFA, as it interacts with other genes through homodimerization, activation, phosphorylation, and dephosphorylation. It also appears to interact with the VEGFA protein via hydrogen bonds. In conclusion, this study suggests that Lenvatinib might be a promising anticancer agent that could treat pheochromocytoma, and VEGF-A, CDC42, PIK3R1, as well as several predicted miRNAs, such as miR-224 and miR-221, might be biomarkers for the early diagnosis.

## INTRODUCTION

According to the National Cancer Institute, in 2022 the number of cancer cases will increase to 1,918,030 and the number of cancer deaths will rise to 609,360 (1)and there are various reasons for the cancer outcome but at the molecular level, the driving genes for cancer growth were found to be around 299, of which 258 are normally found in all types of cancer. In addition, 142 genes were related to at least one type of cancer growth, and 87 genes play a role in at least two types of cancer (2). Pheochromocytoma is a cancer of the adrenal gland, which is also a type of multiple neuroendocrine tumors (NETs) where symptoms are only visible on the affected glands and show excess production of hormones due to the tumor growth (3). Approximately 8 people among 1 million in the world suffer from pheochromocytoma (PHEO or PCC), a rare sporadic neuroendocrine tumor that secretes catecholamines and metanephrines, which cause hypertension and excessive sweating in the patients by altering the epinephrine and norepinephrine hormone production (3). The RET gene mutation is the principal cause of the disorder. Currently, pheochromocytoma is diagnosed with genetic tests, the level of hormones in the blood, and urine. Imaging tests are rarely performed to locate tumors and there is no known cure for the disease except surgical intervention, and no detection technology is suited due to the lack of specificity and sensitivity (3). Thus, it is crucial to investigate the molecular mechanisms involved in pheochromocytoma progression and discover additional biomarkers for effective early diagnosis. In the current scenario, microarray technology has been extensively used to obtain general alterations in the genetic construct during tumorigenesis (4). Bioinformatics strategies are particularly necessary for processing the enormous amount of data generated by microarrays. In this study, the microarray datasets were analyzed to determine differentially expressed genes (DEGs) between diseased and healthy tissues. We used a combination of approaches such as functional enrichment and network analysis together with identifying miRNA-based expression in a particular type of cancer along with their transcription factors to identify significant genes of the Pheochromocytoma.

## Materials and methods

### Data processing

The dataset used for the analysis was the expression profiling of pheochromocytomas performed on the GPL96[HG-U133A] platform. A single gene expression profile (GSE2841) (5) was retrieved from the GEO (Gene expression Omnibus) database, an open-access repository of microarray and next-generation sequencing data (6), which contains 13 extra-adrenal and 62 adrenal tissue tumor samples, as well as one extra-adrenal paraganglioma sample, for 76 samples of adrenal gland tumors (pheochromocytomas). The gene expression data were normalized and annotated with the Affy package (7) and the Limma package of R was used to identify differentially expressed genes between adrenal and extra-adrenal samples (8). Genes that failed to meet the cut-offs of p < 0.05 and log|FC| >1 were omitted from the analysis. The plots for differentially expressed genes were generated via the R package ggplot2 (9).

### PPI network construction and functional pathway enrichment analysis

DAVID Bioinformatics resources is an online database that provides functional annotations for the complete set of genes (10). DEGs are uploaded to the database and analyzed against the human gene ontology map to know the functions of query genes. An overview of the mechanism and cellular processing can be obtained from the interactions between the proteins in this study, PPI networks are created using the search engine Search Tool for Retrieval of Interacting Genes (STRING) (http://string.embl.de) (11) and visualized via Cytoscape (12) with a confidence score of 0.4 as the cut-off criterion. Molecular complex detection (MCODE) is used to screen the modules of the PPI network (13) with a degree cut-off of 2, a score cut-off of 0.2, a K-core equal to 2, and a maximum depth of 100.

### Hub gene prediction

As part of the analysis of the PPI network of differentially expressed genes, we used the CYTOHUBBA plugin, a module of Cytoscape that provides an interface to analyze the topology of protein-protein interaction networks. The score of the subnetwork is calculated from the nodes by Cytoscape to predict the significantly expressed genes (hub genes) (14). Based on the default scoring system of the Cytohubba plugin, the ten top hub genes are ranked which are further used for the analysis.

### Prediction of miRNA and transcription factors and their expression in different types of cancers

The miRNA targets of eight prominent hub genes among the ten in the module were identified through the CSmiRTar (http://cosbi4.ee.ncku.edu.tw/CSmiRTar/search) (15) database that collects information from existing databases such as miRDB, TargetScan, microRNA.org and DIANA-microT (15). For additional information about the expression of miRNAs in particular types of organs, we relied on HMDD v3.2, a database that supports experimental evidence for human microRNAs (miRNAs) and disease association (16). To predict the transcription factors for the top genes in the results of cytohubba, we used a database called TF2DNA (17), which provides evidence of both differentially regulated genes and transcription factor binding motifs. The p-value was adjusted to 0.0001 using the source of the TF2DNA (Computational) method to predict transcriptional factors.

### Homology modeling and molecular docking

#### Homology modeling of proteins

Based on the analysis of the PPI network and hub gene module, we selected the top three genes with more associated connections with other genes in the network. Genes such as CDC42, VEGFA, and PI3KR1 are all primarily involved in cellular signaling pathways and cellular morphology, promote cellular migration, reduce apoptosis, induce endothelial cell division, and are principal components of the signaling process. Further, all three gene products were prepared using homology modeling for the docking studies. All three proteins were modeled by retrieving reference protein structures from the database PBD:2KB0, PDB:3V2A, and PDB:5XGJ, for CDC42, VEGFA, and PI3KR1, respectively (18). Modeller version 9.23 was used to construct the protein 3D structure (19) and the protein model was refined based on E-values and percent identity.

#### Model validation and energy minimization

The RAMPAGE server (http://www-cryst.bioc.cam.ac.uk/rampage) was used to validate the modeled proteins and the Ramachandran plot was constructed by comparing the dihedral ø and ö angles of the amino acid residues (20). ProsA and QMEAN tools (21)are used to determine the packing conformational quality, and Swiss-PDB Viewer simulations are performed to determine the lowest delta G (22).

#### Ligand preparation

A total of 7 lead molecules were obtained from the scientific literature, along with their 3D structures from the drug bank database (23). Furthermore, all molecular properties and biological activities of the ligands were evaluated using Molsoft (24).

#### Study of protein-ligand interactions using auto docking software

The scoring functions of AutoDock software calculate the affinity and interactions between a lead and the target molecules. In this study, we docked all three-target protein molecules against all seven lead or drug compounds by removing water molecules from the target followed by adding polar hydrogens and Kollman charges (25). A rigid grid box was set based on the dimensions of binding residues added via molecular string and the default generic and Lamarckian algorithms were applied to interact with protein and ligand molecules(26). Docked molecules were obtained in different conformations and visualized through a PyMOL viewer and to Ligpolt is used to determine the hydrogen bonds and interactions with the binding sites (27).

#### Molecular dynamic simulation for protein-ligand complex

Molecular dynamic simulations were carried out on the ligand Lenvatinib which is having the best docking score of -8.38 kcal/mol against the VEGFA protein. The MD simulations was performed with the Gromacs 5.1.4 simulation package with Gromos53a6 as a force field. Initially, the system was solvated by adding SPC (Single Point Charge) water. The box type was set to cubic with a starting structure placed at the center at a distance of 1.0 Angstrom (Å) from the wall of the box. The total charge of the system was neutralized by replacing water molecules with corresponding NA+ or CL-counter ions based on the net charge of the system. Then the entire solvated system was energy minimized using the steepest descent algorithm for 50000 steps. During these steps, the Coulombs and Vander Waals interactions were maintained at a cut-off of 0.9 nm and 1.0 nm. Later, the system was equilibrated with a Leap-frog integrator for 100 picoseconds (ps), maintaining simultaneously Berendsen temperature coupling (28) and Parinello-Rahman pressure coupling for 300k and 1 bar, respectively. Additionally, the long-range electrostatics are controlled by Particle-Mesh Ewald (PME) and the short-range electrostatics by Coulombs and Vander Waals at a cut of 1.0nm respectively at a time-step of 10 femtoseconds. LINCS algorithm (29), (30) was used to maintain the geometry of the molecules. Finally, a well-equilibrated system was then subjected to MD simulations for 100 nanoseconds (ns) each at 300 K without any constraints. The conformations were sampled at an interval of 2 ps. The final MD trajectory was analyzed for root mean square deviation (RMSD) of the backbone, root mean square fluctuations (RMSF) of the residues, and also the radius of gyration (g_gyrate) during the production run. The trajectory plots for the above analysis were generated using XMGRACE software (http://plasma-gate.weizmann.ac.il/Grace/).

#### PCA and FEL analysis for sampling the optimal conformations

Principal components (PCs) are the Eigenvectors formed by the projection of points onto the vector, for each such vector, the variance is determined that corresponds to the Eigen value (30). The principal component analysis is a multivariate statistical method that can be applied to address the collective motion of macromolecules concerning their atomic coordinates and the eigenvectors. This method is linearly transforming and reducing the data to a positional covariance matrix (30). The covariance matrix is defined by Ci=< (xi - <xi>) (xj - <xj>)> Where xi is a Cartesian coordinate of the ith Cα atom, N is the number of the Cα atoms considered, and represents the time average over all the configurations obtained. The covariance matrices in this study were generated using the “g_covar” module of GROMACS. The 3N dimensional directions are derived by solving the covariance matrix of the Eigenvectors. In this study, the first 20 Eigenvectors were generated using the “g_anaeig” module, among which only the first two principal components (PC) with minimum deviation and least cosine value of nearly 0.5 were considered for generating Gibb’s free energy (using g_sham script). Further, Free energy landscape analysis (FEL) was performed to sample the various conformations resulting from MD simulation[31]. The PC1 and PC2 were considered as the two reaction coordinates, based on which 2D representation of FEL was generated by implementing the equation: G (q1,q2) = - kB T ln P(q1,q2) where KB is the Boltzmann constant, T is the simulation temperature, and P(q1,q2) is the normalized joint probability distribution (30). Based on this, a 2D projection of the Free energy landscape was generated, and the minimum energy cluster for the macromolecule was determined to identify and also to extract its near-native conformations.

## Results and Discussions

### Identification of DEGs and functional and pathway enrichment

Affy package of R was utilized to normalize and annotate the gene expression data, and LIMMA was used to identify differentially expressed genes between adrenal and extra-adrenal samples (Fig 1). A total of 110 DEGs were obtained based on the criteria applied, i.e., p < 0.05 & log|FC| >1. Among the 97 differentially expressed genes, 43 genes were up-regulated and 54 genes were down-regulated **(S1 Table)** (Fig 2). As pathway enrichment analysis shows both down-regulated and up-regulated genes play key roles in the cell’s biological processes which are all having a main role in cancer progressions such as angiogenesis, cell adhesion, endodermal cell differentiation, MAPK cascade, apoptotic process, cell proliferation, and cell-matrix adhesion contribute to cell proliferation and metastasis pathways crucial to cancer such as the p53 pathway, the PDGF signaling pathway, the interleukin signaling pathway, angiogenesis, apoptosis, the Ras, PI3 kinase, the EGF receptor signaling pathway, the VEGF signaling pathway, the JAK/STAT, Cadherin, and RET signaling pathways (Figs 3-4). Other than pathway enrichment biological and molecular processes concerned with both up and downregulated genes were identified (S1 and S2 Figs).

**Fig 1.**
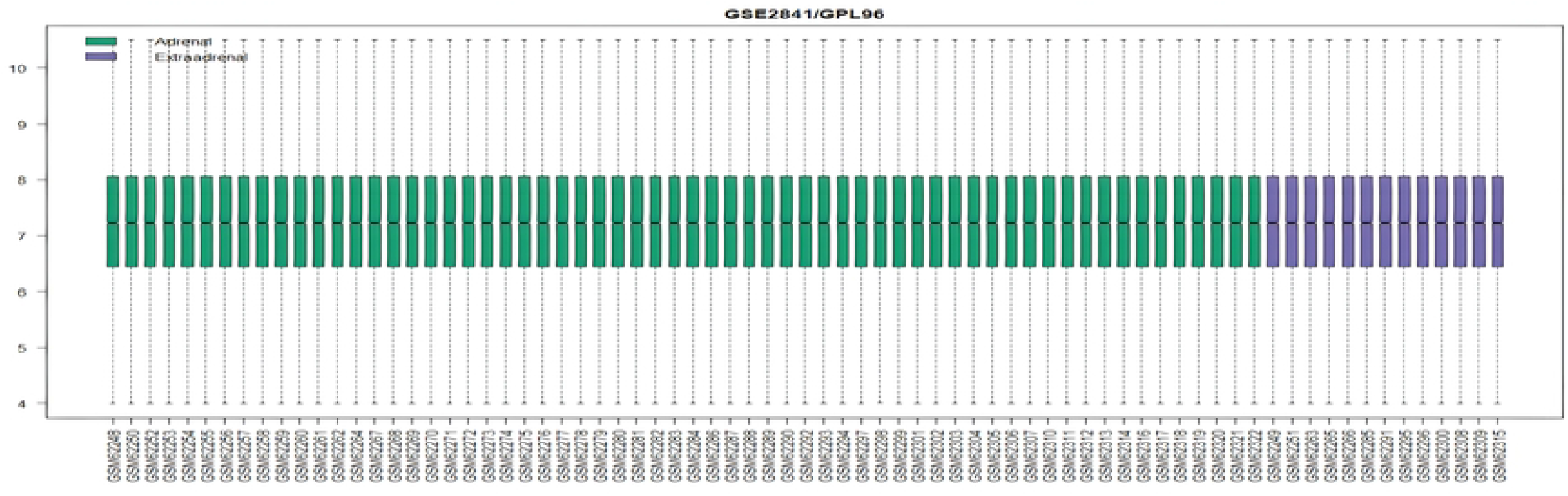
Data Normalization. Microarray data is normalized which indicates the distribution of expression values across the sample size is uniform.

**Fig 2.**
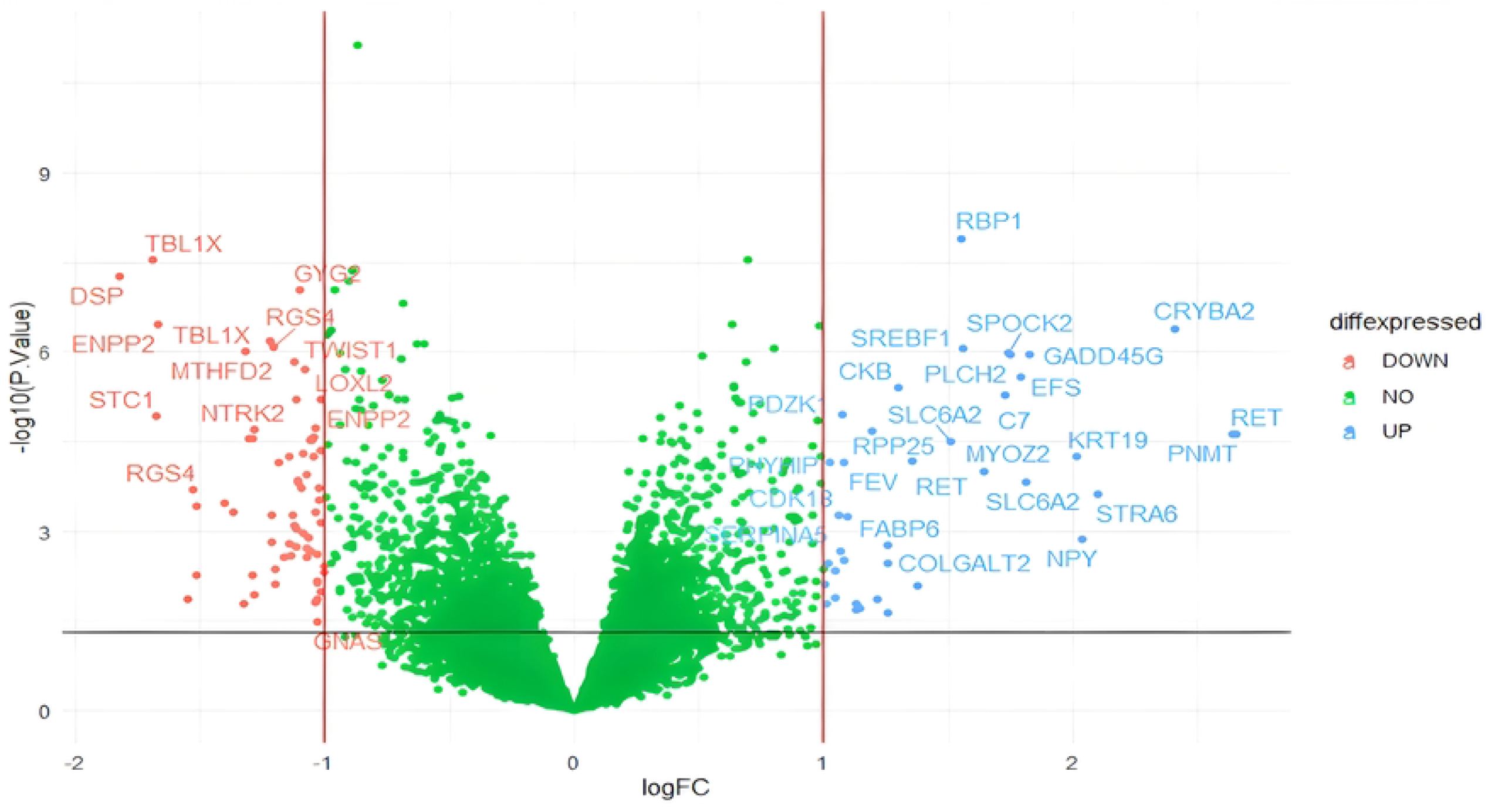
Volcano plot of DEG’s. Cut-off criteria were set at adj. p<0.05 and log|Fc| >1 and genes in blue were upregulated and red are downregulated genes.

**Fig 3.**
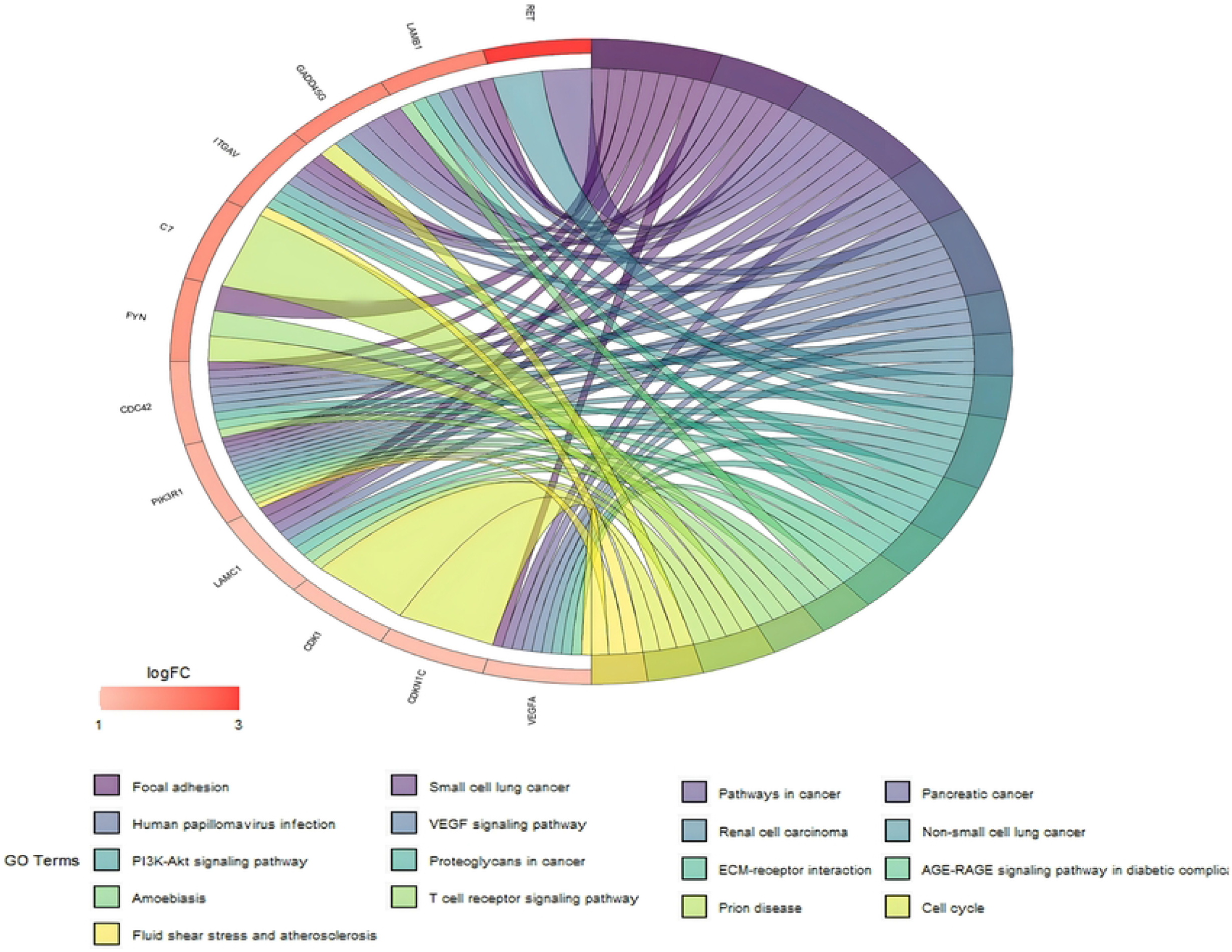
Pathway enrichment of upregulated genes. Genes are functionally enriched by using KEGG pathway terms; Coloured ribbons connect the gene to their associated terms and the intensity of the red color boxes depicts log|Fc| values.

**Fig 4.**
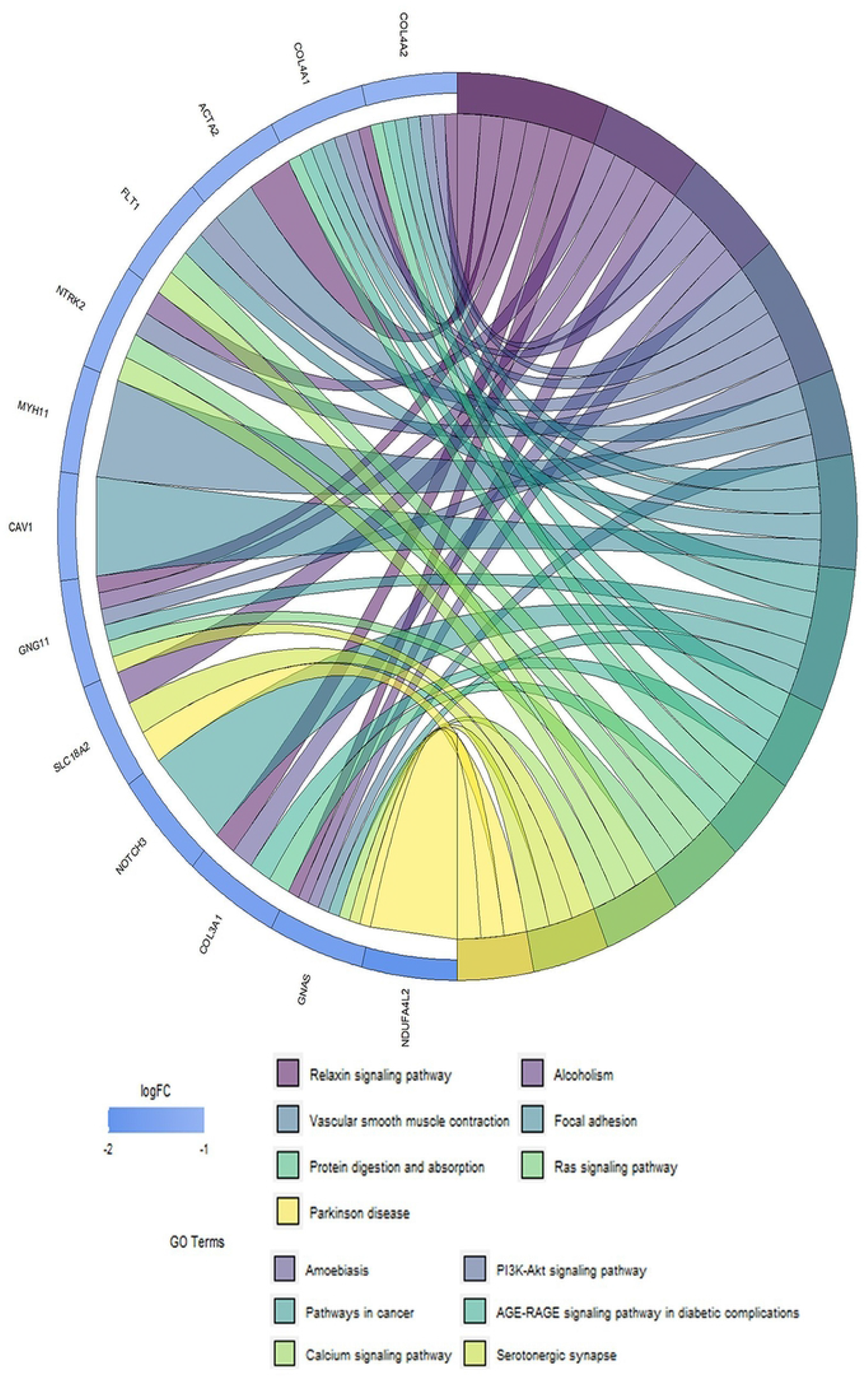
Pathway enrichment of downregulated genes. Genes are functionally enriched by using KEGG pathway terms; Coloured ribbons connect the gene to their associated terms and the intensity of the red color boxes depicts log|Fc| values.

### PPI network construction and hub module predictions

To ensure interactions between DEG’s PPI network are constructed, the STRING database was used to retrieve the protein partners and visualized them with Cytoscape and the network analysis was conducted with the undirected edge network analysis method (Fig 5). A total of eight genes were determined to be hub genes in the network using Cytohubba plugin with a cut-off score of 10; from the network, three significant modules of gene clusters were identified with MCODE with a cut-off score of 5; each module had a different number of nodes and edges.

**Fig 5.**
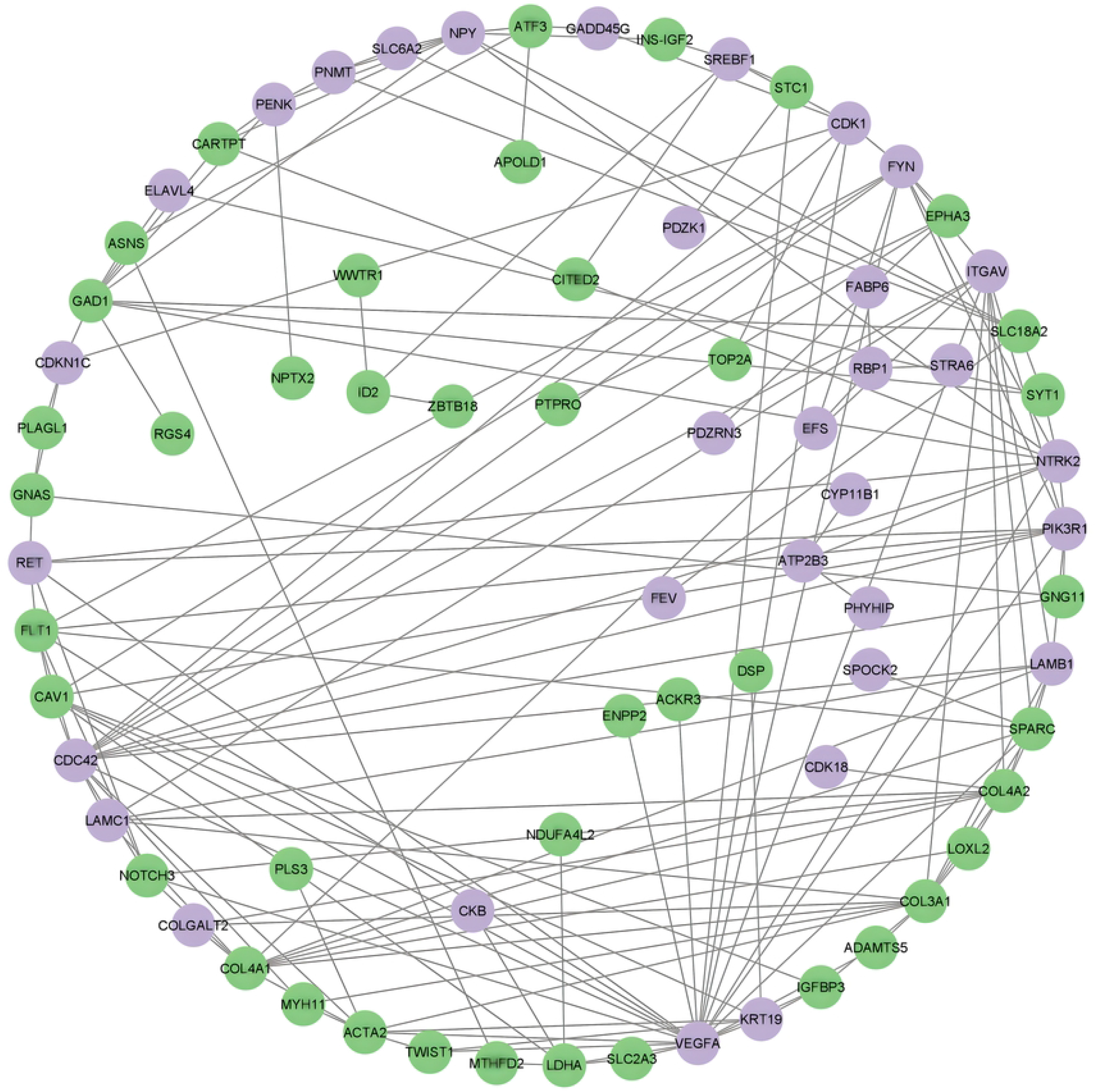
PPI Network. Protein-Protein interaction network analysis of differentially expressed genes (light purple color indicates upregulated genes and green for downregulated genes).

### Functional enrichment Analysis for hub-genes

In order to understand the roles of genes in particular and common pathways for identified top 8 hub genes with a high degree of interconnection within the network, a pathway enrichment analysis was performed (Fig 6). The pathways involved in the JAK-STAT signaling pathway, the PI3K/AKT signaling pathway, the Cell Surface Adhesion pathway, EGFR downregulation, and ECM degradation.

**Fig 6.**
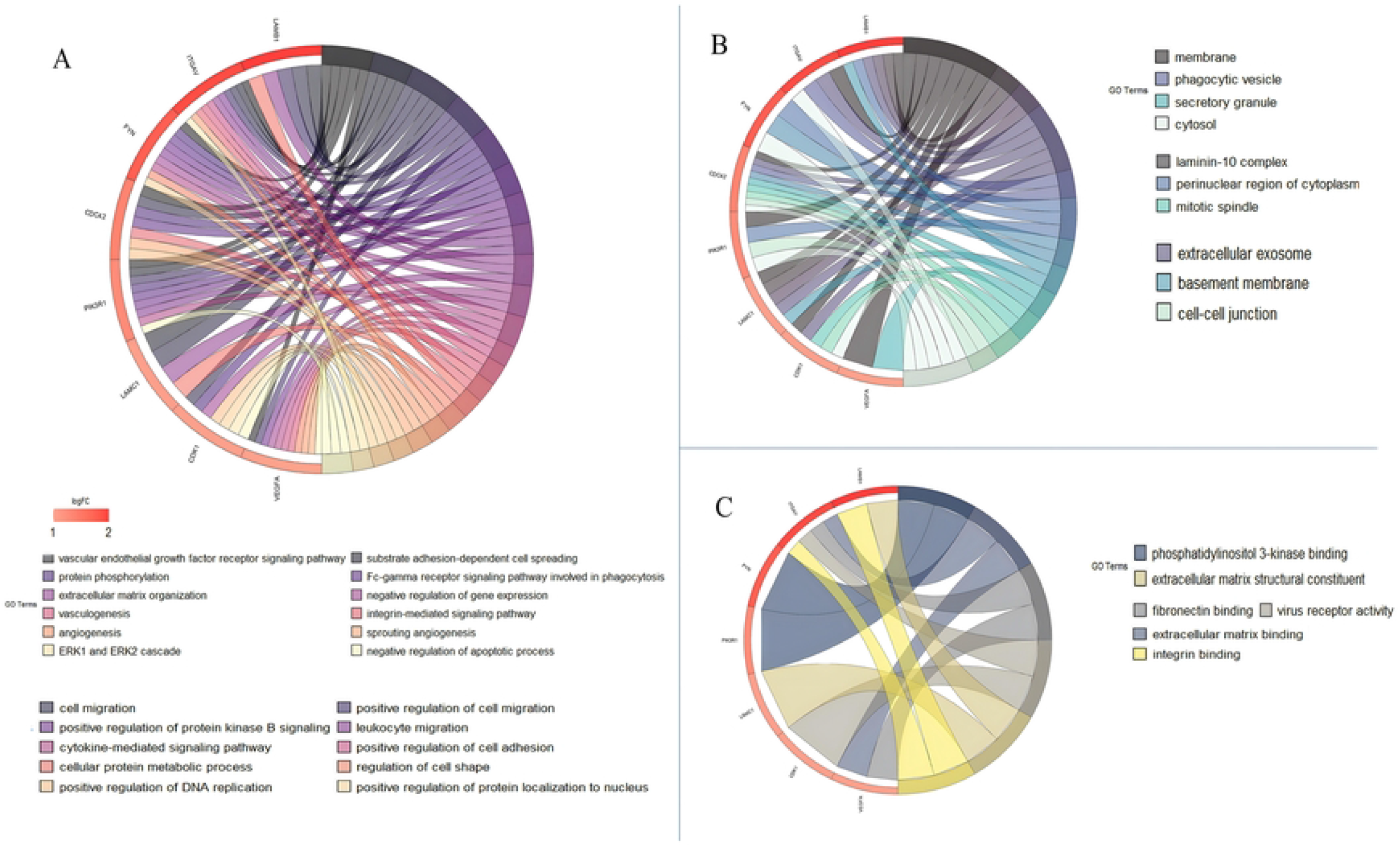
Functional enrichment analysis of hub genes. GO terms associated with hub genes; (A) Biological processes; (B) Cellular Components; (C) Molecular functions.

### Transcription factors and miRNA expression of hub genes

miRNA profiles are predicted for each gene, and their expression in different types of cancer is also validated in (Table 1). Results indicate that miR-5011-5p miRNA concerning the PIK3R1 gene is most significantly upregulated, whereas miR-190a-3p miRNA concerning the VEGFA gene is most significantly downregulated. TF2DNA database is used to determine transcription factors associated with hub genes (Table 2).

**Table 1.** Top Hub genes and their expressions in cancer. Prominent hub genes from the networks are selected and predicted their miRNA targets along with their expressions in different types of cancer.

| GENES | miRNA | Expressed in |
| --- | --- | --- |
| *CDC42* | hsa-miR-214 | Neoplasms of Ovary, Kidney, Thyroid, Cervix, Testicular Germ Cell Tumour |
| *VEGFA* | hsa-miR-200b | Neoplasm of Ovary, Pancreas, Prostate, Carcinomas of thyroid and pancreas, Carcinomas of Endocrine Gland and thyroid |
| *PIK3R1* | hsa-miR-221 | Neoplasms of Ovary, pancreas, Papillary thyroid, Prostate, Renal Cell Carcinomaoma, Carcinomas of Endocrine Gland and thyroid. |
| *ITGAV* | hsa-miR-25 | Neoplasms of pancreas, Ovary, Prostate and carcinomas of papillary thyroid |
| *CDK1* | hsa-miR-31 | Neoplasms of pancreas, Ovary ,Prosta,te and carcinomas of papillary thyroid |
| *LAMB1* | hsa-miR-224 | Prostate Neoplasms, Pancreatic Neoplasms Carcinoma, Renal Cell Ovarian Neoplasms, Thyroid Neoplasms. |
| *LAMC1* | hsa-miR-205 | Neoplasms of ovary, prostate ,pancreas ,thyroid an carcinomas of the prostate and renal cell (clear-cell) and endometrial carcinomas. |
| *FYN* | hsa-miR-96 | Pancreatic Neoplasms Carcinoma, Prostate Prostate Neoplasms Carcinoma, Renal Cell, Clear-Cell Carcinoma, Ovarian |

**Table 2.**
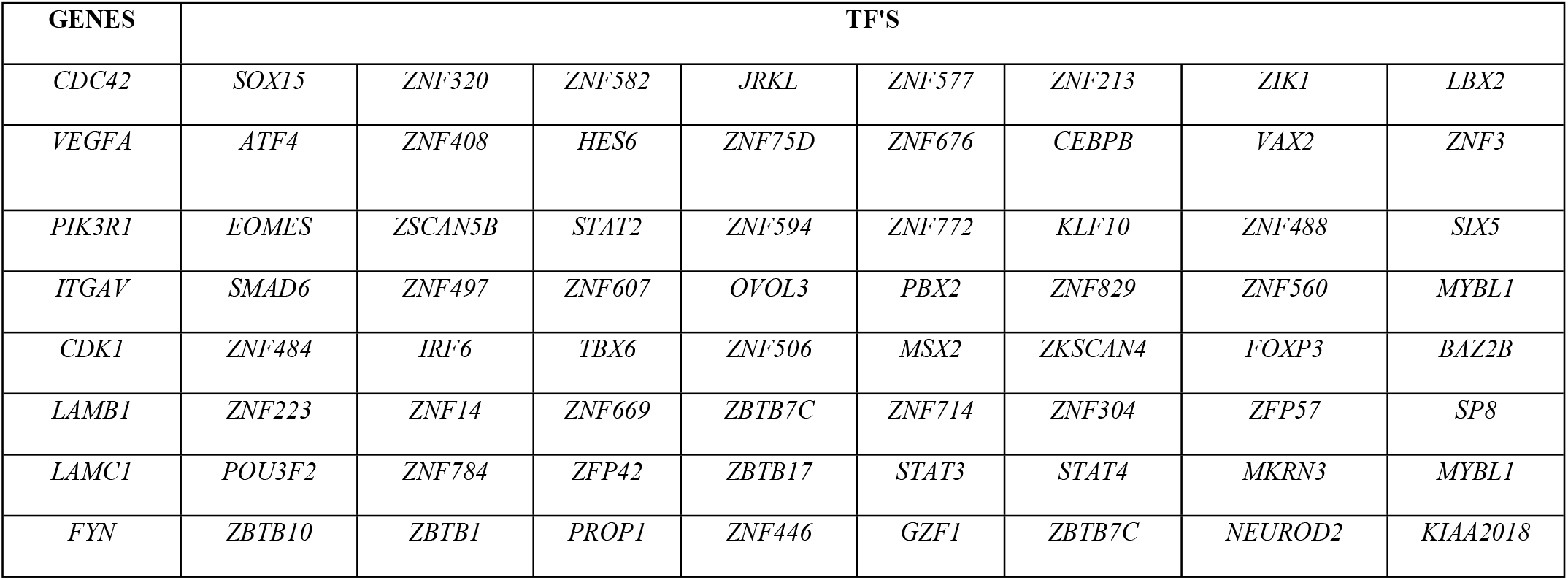
Genes and their TFs. Top hub genes from the network are subjected to predict their transcription factors and major TFs are listed in this table.

### Homology modelling and validation

Modeller 9.23 was used to construct 3-D models of the CDC42, VEGFA, and PI3KRI, and Ramachandran plot analysis to evaluate the stereochemical properties of the modeled proteins. The plot of ø and ö revealed that the homology model of CDC42 has 98.9% residues in the favored region and 0.6% of residues in the allowed region as well as the outlier region (S3 Fig). Modeled VEGFA results in 98.9% in the favored region 1.1% in the allowed region and 0% in the outlier region (S3 Fig) and PI3KRI 99% in the favored region, 1% in the allowed region, and 0% in the outlier region (S3 Fig). The overall quality of the 3-D modeled proteins was measured by ProSA and QMEAN Servers. Z scores from the ProSA server for all models of proteins were -7.67, -3.62, and -6.04 respectively and QMEAN4 scores for all 3 modeled proteins were -0.07, -0.98, and -0.54 respectively.

### Properties of ligands

We evaluated lead molecules obtained from the literature using a variety of tools to determine their molecular properties and bioactivity, and seven molecules were considered for virtual screening and Lipinski’s rule of drug and compound bioactivity was taken into consideration. In the case of Sorafenib, Lenvatinib, Vandetanib, and Sunitinib, they all obey Lipinski’s rule, and their bioactivity is appreciable as well **(Tables 3 and 4)**.

**Table 3.** Molecular properties of ligand molecules. Molecular properties of ligand molecules are predicted in order to full fill the Lipinski rule and the molecules such as Cabozantinib, Lanreotide, and Everolimus show violations thus expect these ligands’ rest were selected for docking studies.

| Drug name | no of heavy atoms | mo. Weight gm/mol | hydrogen acceptors | hydrogen bond donors | no of violations | rotatable bonds | Drug likeness |
| --- | --- | --- | --- | --- | --- | --- | --- |
| Sorafenib (31) | 32 | 464.82 | 7 | 3 | 0 | 9 | 0.55 |
| Lenvatinib (32) | 30 | 426.85 | 5 | 3 | 0 | 8 | 0.55 |
| Vandetanib (33) | 30 | 475.35 | 6 | 1 | 0 | 6 | 0.55 |
| Cabozantinib (34) | 37 | 501.51 | 7 | 2 | 1 | 10 | 0.55 |
| Lanreotide (35) | 77 | 1096.32 | 12 | 13 | 3 | 19 | 0.17 |
| Sunitinib (36) | 29 | 398.47 | 4 | 3 | 0 | 8 | 0.55 |
| Everolimus (37) | 68 | 204.66 | 14 | 3 | 2 | 9 | 0.17 |

**Table 4.** Predicted biological properties of ligand molecules. Molecular properties of ligand molecules are predicted in order to full fill the Lipinski rule and the molecules such as Cabozantinib, Lanreotide, and Everolimus show violations thus expect these ligands’ rest were selected for docking studies.

| Drug name | GPCR<br>ligand | Ion channel<br>modulator | Kinase<br>inhibitor | Nuclear receptor<br>ligand | Protease<br>inhibitor | Enzyme<br>inhibitor |
| --- | --- | --- | --- | --- | --- | --- |
| Sorafenib (31) | 0.18 | 0 | 0.44 | -0.07 | 0.11 | 0.08 |
| Lenvatinib (32) | 0.18 | 0 | 0.62 | -0.23 | 0.14 | 0.16 |
| Vandetanib (33) | 0.05 | -0.12 | 0.64 | -0.45 | 0.29 | 0.04 |
| Cabozantinib (34) | 0.06 | -0.1 | 0.43 | -0.01 | 0.07 | 0.05 |
| Lanreotide (35) | -3.66 | -3.82 | -3.8 | -3.82 | -3.55 | -3.72 |
| Sunitinib (36) | -0.16 | -0.62 | 0.51 | -0.8 | -0.51 | -0.23 |
| Everolimus (37) | -3.36 | -3.61 | -3.68 | -3.59 | -2.97 | -3.16 |

### Molecular docking studies

Molecular docking studies are conducted to determine whether the ligand-target complex can bind with the highest affinity between their active sites or not, which are then applied to molecular dynamics simulations to determine whether the interaction of a small molecule with a protein leads to its inactivation or activation. Autodock tools V.1.5.6 were used for docking and Swiss PDB viewer for energy minimization of all proteins (22) while the docked results are visualized with PyMOL, and interactions are examined with LIGPLOT (27). The docking is performed for CDC42, VEGFA, and PI3KRI proteins individually, and results are analyzed based on binding energy scores and interactions between molecules.

In the case of CDC42 predicted active sites are used to perform rigid docking, and the results indicated that Sorafenib has better binding energy of - 7.05 kcal/mol along with four hydrogen bond interactions. Whereas the docking results of VEGFA and PI3KRI show that Lenvatinib has the best binding energy score, -8.38 (Fig 7) and -8.07 kcal/mol respectively, with the variation in hydrogen bond interactions i.e. two interactions between VEGFA and Lenvatinib and no hydrogen bonds interactions between PI3KRI and Lenvatinib (Table.5)

**Table 5.** Molecular docking studies. The drugs which full fills the Lipinski rules are further used for docking studies in which the complex with the best binding energy is further used for molecular dynamics simulation studies.

| Target: Ligand | Interacting Residues | n.o/h-bonds | Binding energy (kcal/mol) |
| --- | --- | --- | --- |
| CDC42: Sorafenib | Thr 17,Glu31,Pro34 | 4 | -7.05 |
| CDC42:Lenvatinib | Glu31 | 0 | -5.66 |
| CDC42:Vandetanib | Pro34 | 0 | -5.67 |
| CDC42:Sunitinib | Val33, Glu31 | 2 | -5.93 |
| VEGFA: Sorafenib | Cys47,Glu51,Tyr8 | 3 | -7.33 |
| VEGFA:Lenvatinib | Cys47,Asn49 | 2 | -8.38 |
| VEGFA: Vandetanib | Asn49 | 1 | -7.77 |
| VEGFA: Sunitinib | Cys47 | 2 | -8.31 |
| PI3KRI: Sorafenib | Asn49 | 0 | -3.58 |
| PI3KRI:Lenvatinib | Leu80, Phe68, Gly67, Phe79, Tyr66, Ser75, Val78,Ser76 | 0 | -8.07 |
| PI3KRI:Vandetanib | Val33,Glu31 | 0 | -6.53 |
| PI3KRI: Sunitinib | Cys47, Glu51 | 2 | -4.23 |

**Fig 7.**
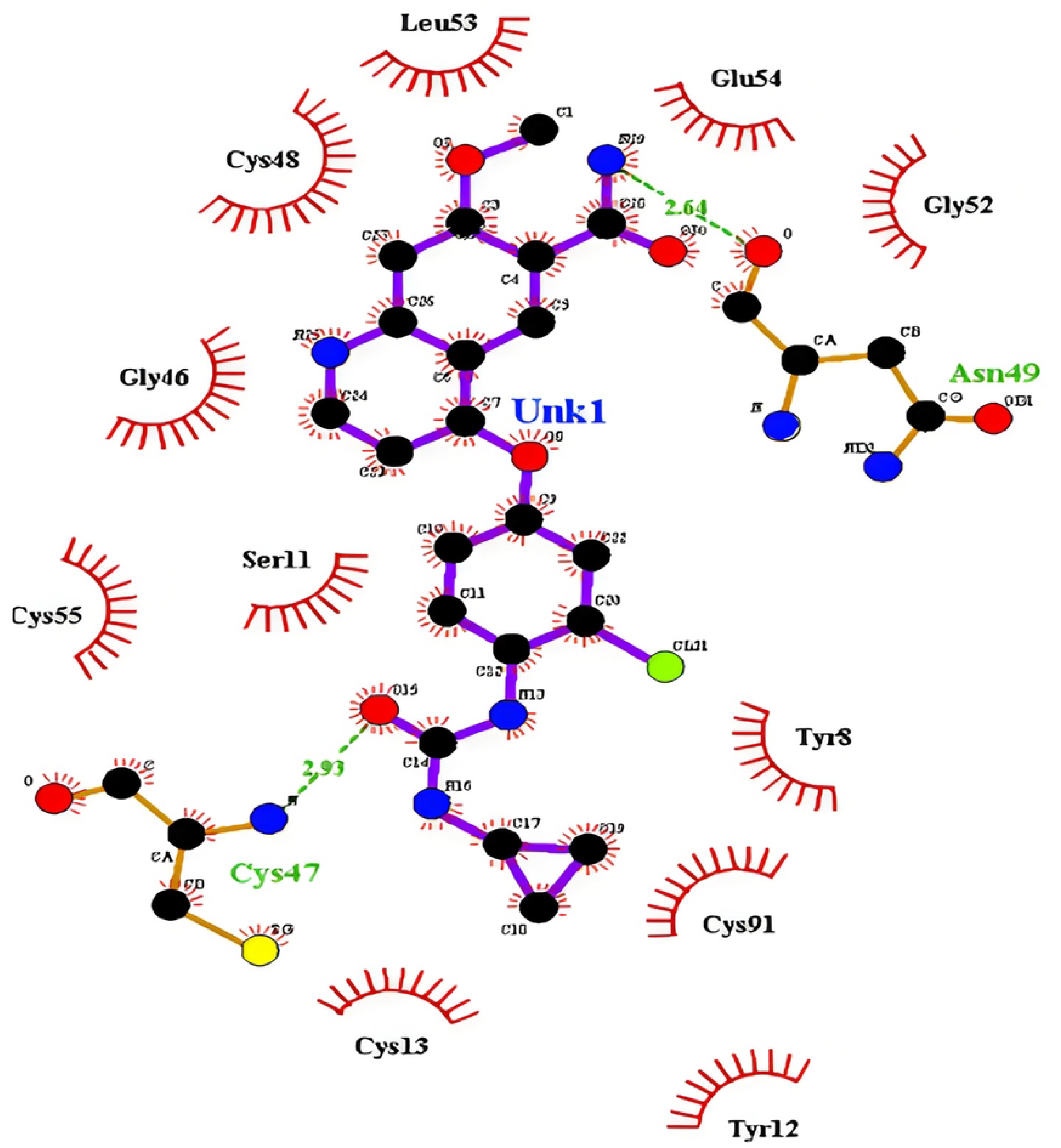
Docking result. The interactions between VEGFA and Lenvatinib, green colored amino acid shows hydrogen bond interaction with the protein.

### Molecular dynamics simulation analysis

The RMSD (Root Mean Square Deviation) profile (Fig 8A) of the complex confirms that the complex has equilibrated after 40 ns with an RMS deviation of about ∼0.5-0.62 nm. Other than the N-terminal residues, the residues Y26, D50, and Q74 have undergone higher fluctuations of about ∼0.45-0.40 nm as shown in the RMSF plot (Fig 8B). Initially, the complex has shown compactness, however, it reached a higher Rg value of ∼1.9nm thereby attaining open conformation after 45ns (Fig 8C). This highly correlates with the SASA plot (Fig 8D), which indicates increased solvent accessibility ranging from about 70-80nm^2^. The stability of the protein-ligand complex was analyzed based on the inter hydrogen bonds and it was noted that the number of hydrogen bonds varied from 4-to 6, however, 5 inter hydrogen bonds are stably maintained between the protein and ligand throughout the MD production run, thereby indicating increased stability of the complex (Fig 9A). Moreover, the minimum potential energy structure of the complex (Fig 9C) was attained at 27.29ns with a minimum energy of -8.4e+05 (kJ/mol) as shown in the potential energy plot (Fig 9B).

**Fig 8.**
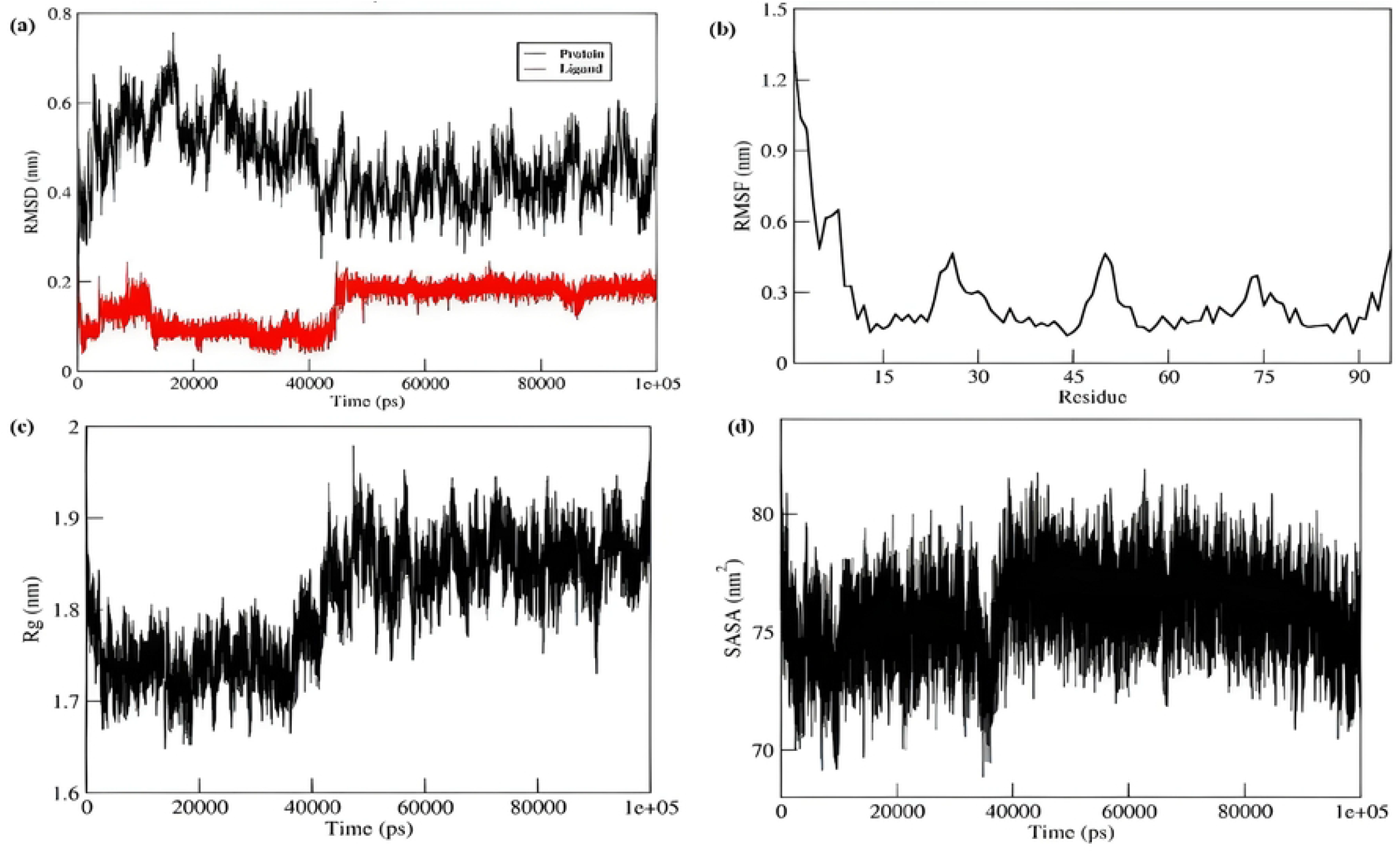
MD simulation trajectories’ analysis. **(A)** RMSD plot showing the that the complex has equilibrated after 40 ns. **(B)** RMSF plot shows the fluctuations about ∼0.45-0.40 nm in residues of Y26, D50 and Q74. **(C)** Radius of gyration plot Initially, the complex has shown compactness, however, it reached a higher Rg value of ∼1.9nm thereby attaining open conformation after 45ns. **(D)** Solvent Accessible surface area.

**Fig 9.**
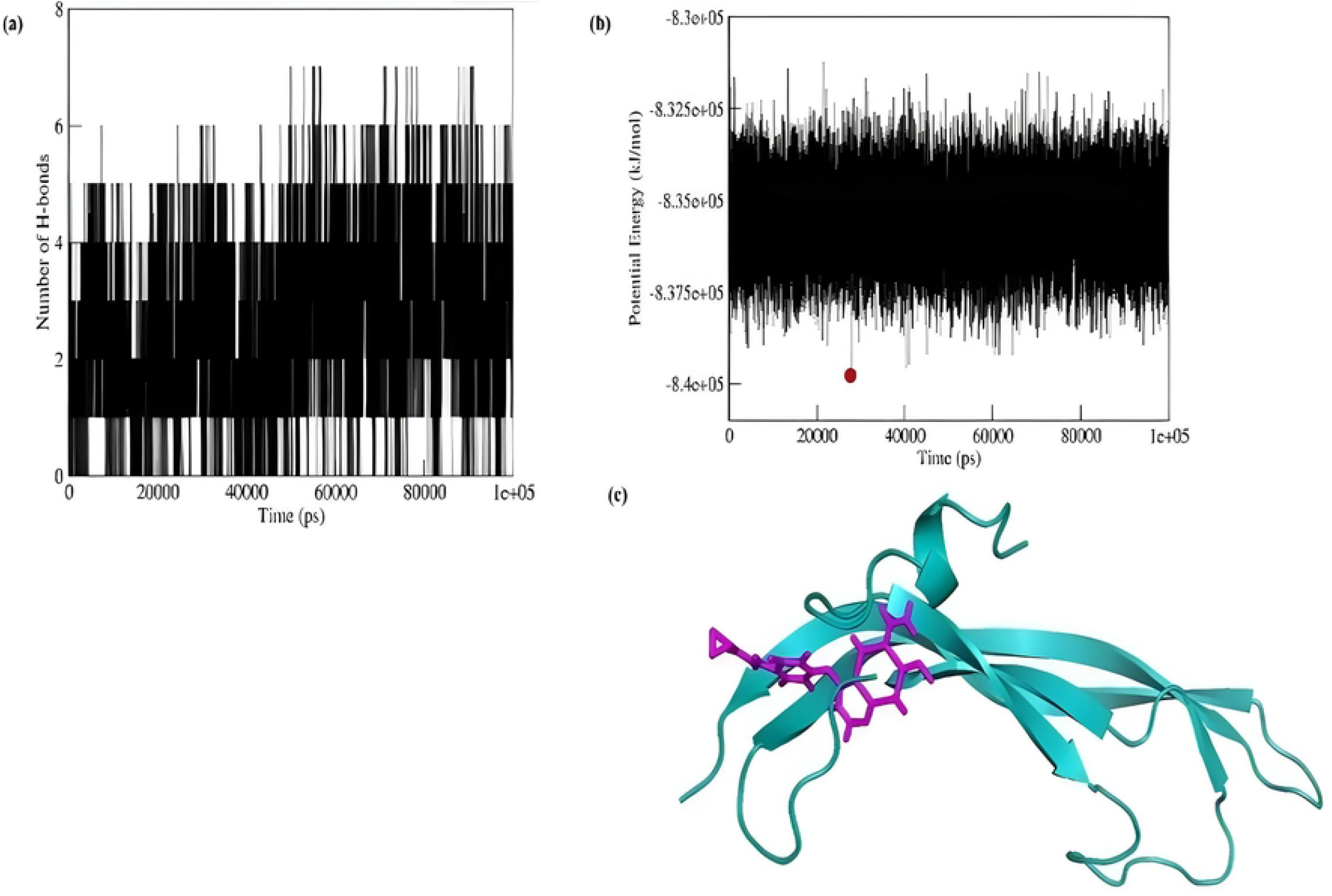
Interactions between protein and ligand. **(A)** Number of hydrogen bonds between protein and ligand throughout the MD production run, **(B)** Potential Energy Plot, **(C)** Minimum potential Energy Structure obtained at 27.79 ns.

In this study, Principal component analysis (PCA) was implemented to identify optimal conformations along each trajectory. Therefore, eigenvectors 2 and 3 with minimum deviation and least cosine value of nearly ∼0.5 were considered as two principal components (PC) for generating a 2D projection of the Free energy landscape (Fig 10A) This resulted in a single minimum of near-native conformation, which denotes the stable conformation of the complex. The representative structure of the near-native conformation (Fig 10B) was obtained at the coordinate (−0.03228,-0.02462) which falls at the time frame of 51958^th^ ps. On comparative analysis of the Initial and near-native conformation of the complex, it was inferred that the α helix (Y26-E31) and the loop region (K71-Q76) have been transformed as loop region and β sheet respectively in the near-native complex as shown in (Fig 11) and **Table 6**.

**Table 6.** Comparative interaction analysis of the initial and the near-native conformation of the complex

| Protein-Ligand complex | Number of interactions within the complex |  | Residues interacting with ligand |  |
| --- | --- | --- | --- | --- |
|  | Hydrophobic Interactions | Hydrogen Bonds | Hydrophobic Interactions | Hydrogen bonds |
| Initial complex | 3 | 4 | Tyr8, Leu53 | Cys13, Cys47, Asn49, Gly52 |
| FEL near-native complex | 3 | 6 | Gly46, Leu53 | Ser11, GLy46, Asn49, ASp50 |

**Fig 10.**
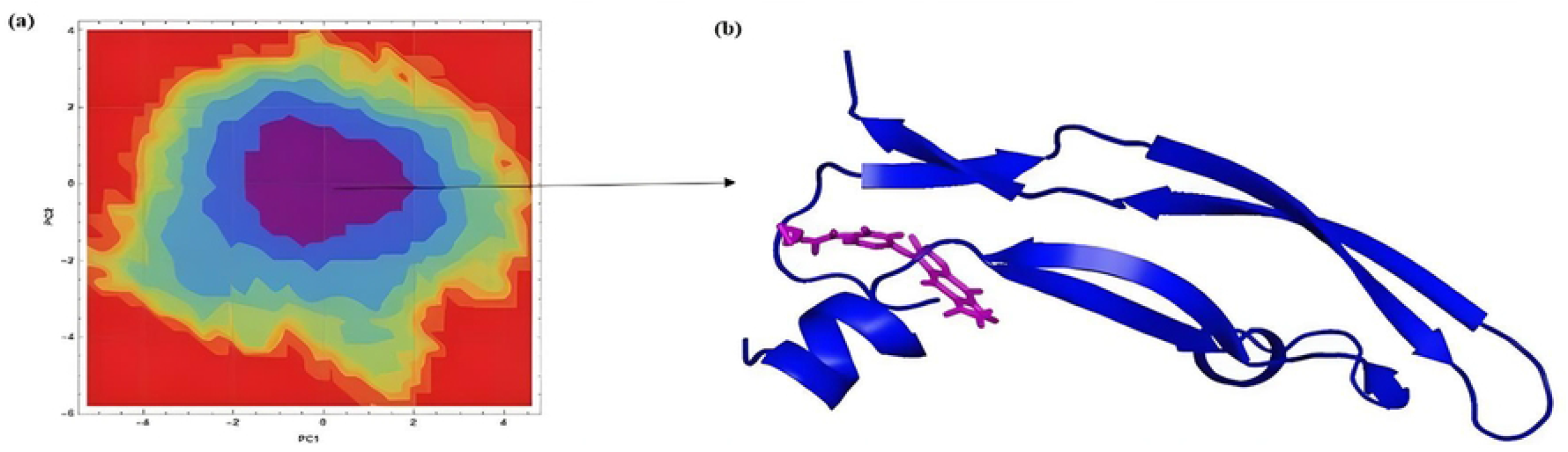
Free Energy landscape analysis. **(A)** 2D projection of the free energy landscape contour map of HPSE at pH 7.0, **(B)** Corresponding minimum energy cluster structure.

**Fig 11.**
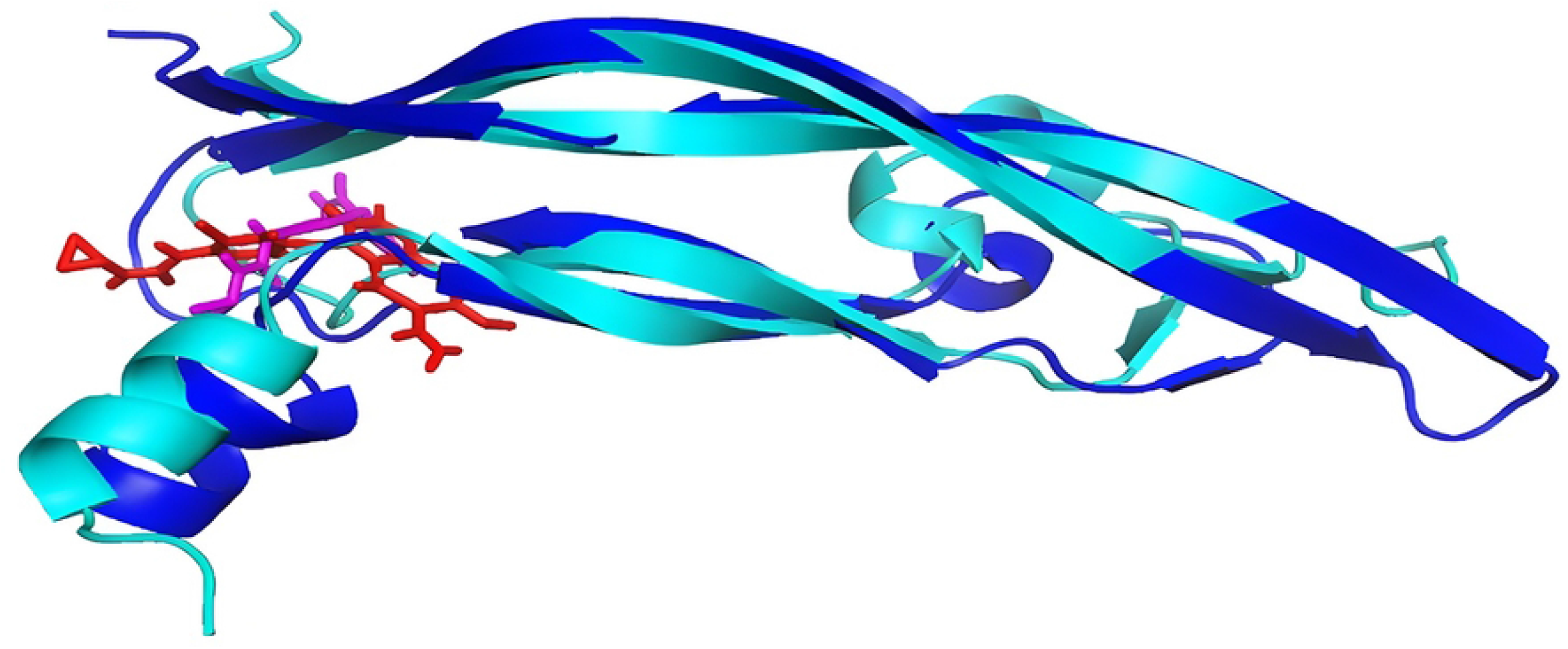
Comparative analysis. The initial (Cyan) and near-native conformation (Blue), where the protein was represented as cartoon and ligand as sticks.

## Discussions

This study considered both up-and-down regulated genes and identified both DEGs are playing role in pathways and processes which are involved with cancer cells. The major pathways and processes include PDGF signaling, Interleukin signaling, Angiogenesis, Apoptosis, EGF receptor signaling, VGF signaling, JAK/STAT, cadherin, and RET, p53, Ras, PI3 kinase pathway. In this study, we found that the CDC42, VEGFA, and PIK3R1 genes were highly expressed with multiple interactions in the protein-protein network, suggesting a prominent role in the progression of pheochromocytoma. As an integral part of the cytoskeletal arrangements, CDC42 manages cell-to-cell adhesions, which belong to the Rho family and are critical for regulating cell morphology, migration, and cell cycle progression (45). Increased expression of CDC42 has been observed in many different types of cancer including Ovarian Neoplasms (46),Kidney Neoplasms, Thyroid carcinoma (47), Cervical Neoplasms, and Testicular Germ Cell Tumours. A CDC42-activated kinase promotes the adhesion and invasion of cells by activating PAK1 and PAK2, which are associated with p21-activated kinase (48). A variation in the expression levels of CDC42 could directly influence cell proliferation and suppress apoptosis in cancer cells. VEGFA encodes a heparin-binding protein that exists as a homodimer with di-sulfide bondage that acts as a growth factor involved in angiogenesis (adult stage), vasculogenesis (embryonic stage), and endothelial cell growth and proliferation along with cellular migration (47). This complex of VEGF-A and VGFR2 dimer and activated CDC42 dephosphorylates MAPK38 into Phospho-MAPKinaseP38, which is the main precursor of the RAF/MAPK cascade and results in cellular proliferation. It is found that the expression of VEGF-A is altered in ovarian neoplasms, thyroid neoplasms, thyroid carcinomas, ovarian endometrial neoplasms, pancreatic neoplasms, prostate neoplasms, adenocarcinoma, pancreatic carcinoma, and others (49). Homodimerization of VEGF results in the formation of VEGF-A, VEGF-B, VEGF-C, and VEGF-D. The receptor VEGFR2 dimerizes and is activated with VEGF-A dimer, which initiates the downstream cascade, involving CDC42, which triggers the p38 MAPK pathway. This complex of VEGF-A and VGFR2 dimer and activated CDC42 dephosphorylates MAPK38 into Phospho-MAPKinaseP38 which is the main precursor of the RAF/MAPK cascade and results in cellular proliferation. It is found that the expression of VEGF-A is altered in ovarian neoplasms, thyroid neoplasms, thyroid carcinomas, ovarian endometrial neoplasms, pancreatic neoplasms, prostate neoplasms, adenocarcinoma, pancreatic carcinoma, and others (49). The PIK3R1 gene encodes a protein that binds to activated protein-Tyr kinases and acts as an associate protein of the p110 catalytic unit of the plasma membrane (50). PIK3R1 expression is altered in most types of neoplasms, including prostate, renal, thyroid, pancreatic, and ovarian carcinomas. It is majorly involved in the cell signaling response to FGFR1 and VEGFA-VEGR2 dimer complexes, as well as major pathways for cell proliferation and migration. The PIK3R1 is a subunit of the PI3K that binds with the VEGFA-VEGF2 dimer to activate the PIP2 to PIP3, which in turn activates the AKT signaling pathway. It is an important pathway in cancer progression. So, the three hub genes have been identified as playing a key role in VEGF signaling (signal transduction pathways, which are responsible for cell proliferation and migration). Targeting the VEGFA pathway in the case of pheochromocytoma may be beneficial in treating patients with this cancer, thus this study examines the drugs that are currently in clinical trials to treat MEN (Multiple endocrine neoplasia) since pheochromocytoma is also a type of endocrine gland-related cancer. These compounds are predicted as potential drugs to treat Pheochromocytoma. Results from simulations and FEL studies support the hypothesis that Lenvatinib (currently in phase III trial) inhibits VEGFA overexpression in pheochromocytoma patients. Hence, Lenvatinib may be the most effective drug to target VEGFA in patients with pheochromocytoma. This study also identified miRNAs associated with the hub genes and miR-244 is the most significantly upregulated miRNA, whereas miR-221 is the most significantly downregulated miRNA. As miR-221 is also related to the progression of endocrine gland carcinomas and prostate cancers and high levels of miR-224 are positively associated with the progression of neoplasms of the thyroid, ovarian, and pancreatic glands, it could serve as a biomarker for the diagnosis of the Pheochromocytoma.

## Conclusions

Despite advancements in medical technologies, the difficulty in early detection and the lack of effective treatments in advanced stages have made it very difficult to diagnose and treat pheochromocytoma patients. So understanding the etiological factors and molecular mechanisms is very crucial for effective treatment. As a result of this study, we have discovered that differentially expressed genes within the interactome, as well as highly interacting hub genes, can be used as biomarkers in conjunction with their respective miRNAs and TRFs. VEGFA gene interacts with other genes by homodimerizing, activating, phosphorylating, and dephosphorylating most major components of cellular pathways (Fig 12), and in a study targeting VEGFA as a drug target, this gene showed the strongest affinities with Lenvatinib. Through the interaction analysis, it was revealed that the stable interactions of Lenvatinib with the VEGFA protein appeared to be mediated by H-bonds. Thus, this study suggests that Lenvatinib is considered to be a potentially effective anticancer agent to treat pheochromocytoma and VEGF-A, CDC42, PIK3R1, and several predicted miRNAs such as miR-224, miR-221 might be key biomarkers related to pheochromocytoma.

**Fig 12.**
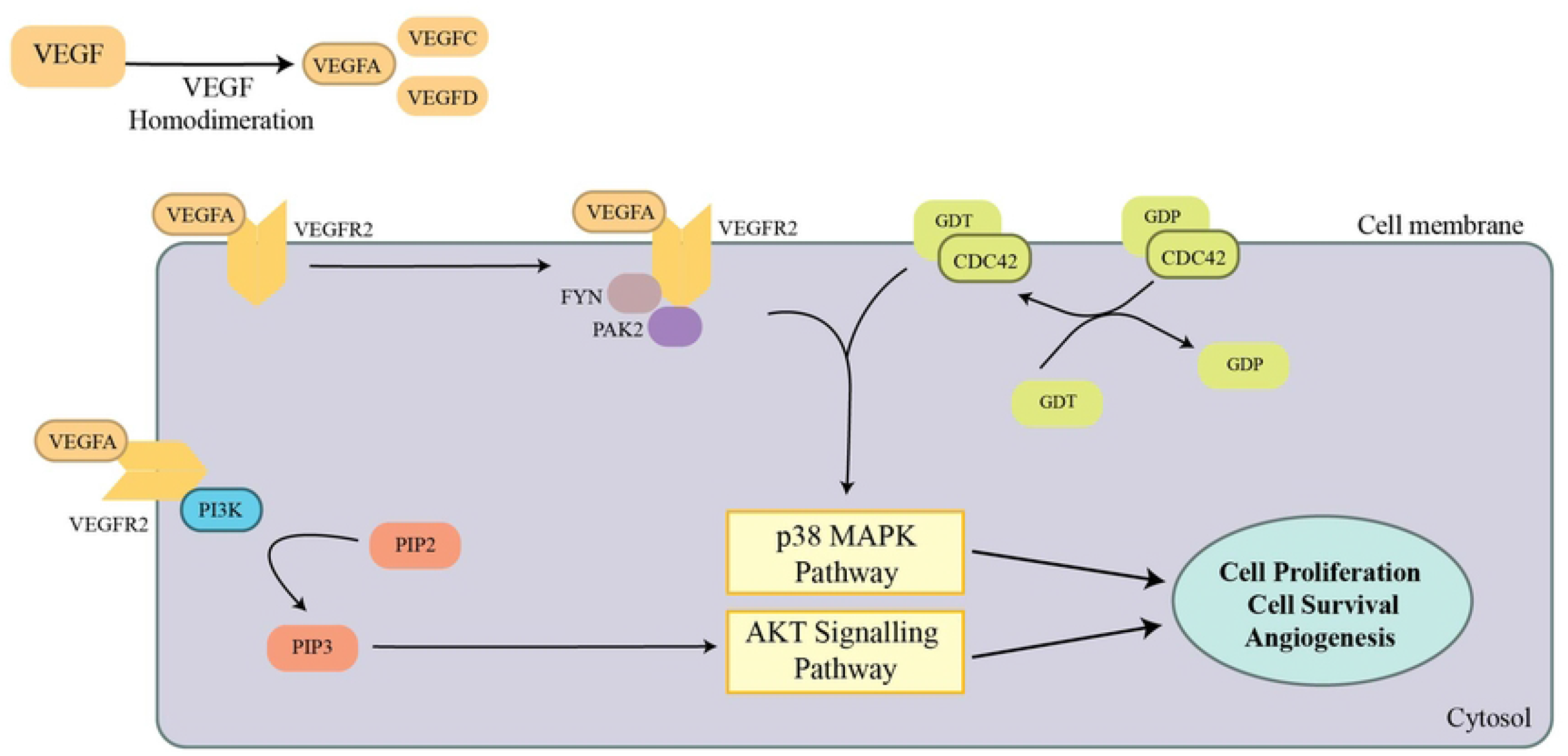
The Central role of VEGFA in cancer outcome. The VEGFA signaling pathway determines how targeted hub genes (VEGFA, CDC42, PI3K) are having a role in cancer-causing signaling pathways such as p38 MAPK and AKT signalling pathways.

## Contributions of authors

**MA** conceptualization, methodology and designed the study, supervised by **JJ**, investigated by **CC**, Data Curation, Formal Analysis, Visualization, and Writing-Original Draft preparation are contributed by **SPJ**.

## Limitations

Our work doesn’t validate with the live samples and it only enlights the possible biomarkers along with the best drug only with *Insilco* studies.

## Acknowledgment

We greatly acknowledge Department of Science and Technology (F.No. EMR/2016/000498 dated: 26.09.2016), TANSCHE (File No. RGP/2019-20/ALU/HECP-0049), and DST Indo-Taiwan (GITA/DST/TWN/P-86/2019) for providing financial assistance through Major research projects. In addition, we acknowledge the Department of Biotechnology, Bioinformatics and Computational Biology Centre (DBT-BIC). BT/PR40154/BTIS/137/34/2021, DST-FIST (SR/FST/LSI-667/2016), New Delhi for providing infrastructure facilities. We sincerely acknowledge the Department of Science and Technology, New Delhi for the financial support in general and infrastructure facilities sponsored under PURSE 2nd Phase programme (Order No. SR/ PURSE Phase 2/38 (G) dated: 21.02.2017) and MHRD-RUSA 2.0 (F.24-51/2014-U, Policy (TNMulti-Gen), Dept. of Edn. Govt. Of India, Dt.09.10.2018. In addition to thanking VIT University for providing the computational facilities, we are also grateful to Mr. Shogun Sugumar for his assistance in analysing data obtained from the microarrays.

## Supporting Information

**S1 Fig. Over-representation of analysis using GO terms of the up-regulated genes. (A)** Biological processes, **(B)** Molecular Functions, **(C)** Cellular Processes.

**S2 Fig. Over-representation of analysis using GO terms of the down-regulated genes. (A)** Biological processes, **(B)** Molecular Functions, **(C)** Cellular Processes.

**S3 Fig. Ramachandran plot Validations. (A)** Modeled CDC42, **(B)** Modelled VEGFA, **(C)** Modelled PIK3KRI.

**S1 Table. Differentially expressed genes in the dataset after normalization**.

## GRAPHICAL ABSTRACT

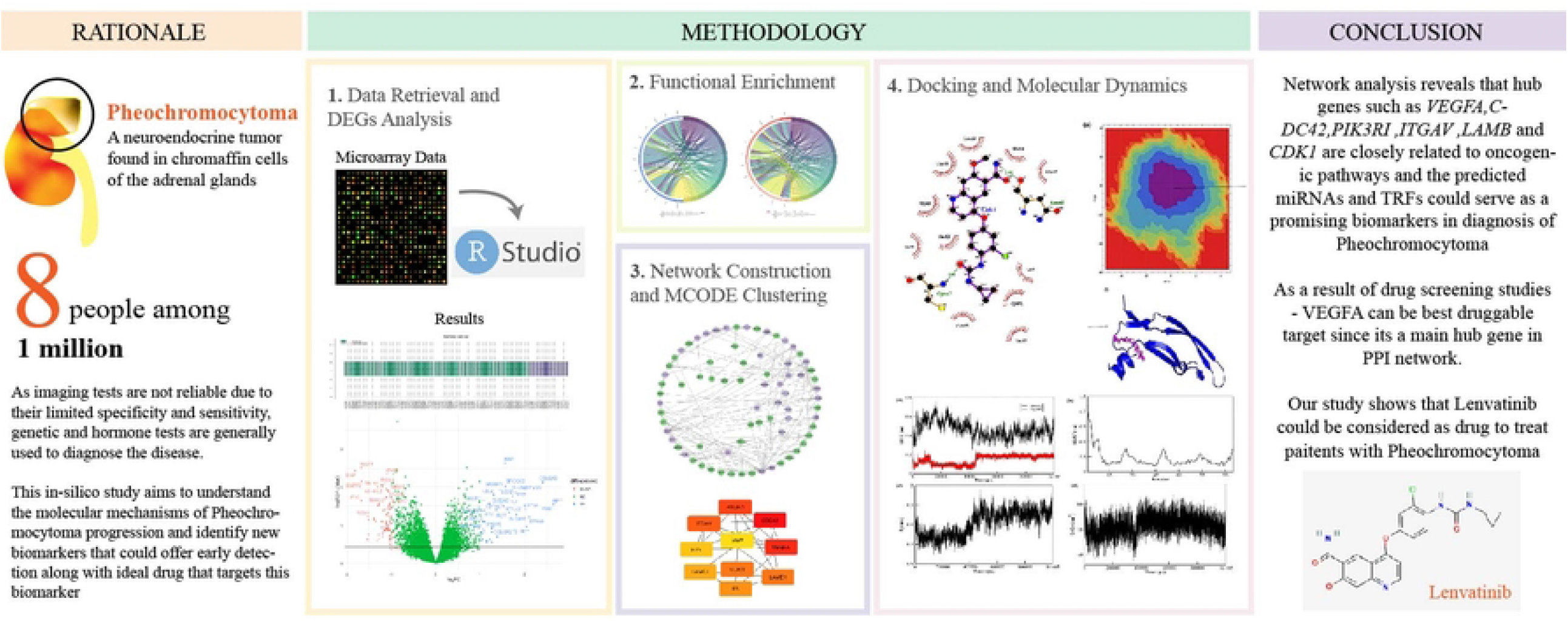

